# Histology-associated transcriptomic heterogeneity in ovarian folliculogenesis revealed by quantitative single-cell RNA-sequencing for tissue sections with DRaqL

**DOI:** 10.1101/2022.12.14.520513

**Authors:** Hiroki Ikeda, Shintaro Miyao, So Nagaoka, Takuya Yamamoto, Kazuki Kurimoto

## Abstract

High-quality single-cell RNA-sequencing (RNA-seq) with spatial resolution remains challenging. Laser capture microdissection (LCM) is a widely used, potent approach to isolate arbitrarily targeted cells from tissue sections for comprehensive transcriptomics. Here, we developed DRaqL (direct RNA recovery and quenching for LCM), an experimental approach for efficient lysis of single cells isolated by LCM from alcohol- and formalin-fixed sections without RNA purification. Single-cell RNA-seq combined with DRaqL allowed transcriptomic profiling from alcohol-fixed sections with efficiency comparable to that of profiling from freshly dissociated cells, together with effective exon– exon junction profiling. Furthermore, the combination of DRaqL and protease treatment enabled robust and efficient single-cell transcriptome analysis from tissue sections strongly fixed with formalin. Applying this method to mouse ovarian sections, we revealed a transcriptomic continuum of growing oocytes quantitatively associated with oocyte size, and detected oocyte-specific splice isoforms. In addition, our statistical model revealed heterogeneity of the relationship between the transcriptome of oocytes and their size, resulting in identification of a size–transcriptome relationship anomaly in a subset of oocytes. Finally, we identified genes that were differentially expressed in granulosa cells in association with the histological affiliations of granulosa cells to the oocytes, suggesting distinct epigenetic regulations and cell-cycle activities governing the germ–soma relationship. Thus, we developed a versatile, efficient approach for robust single-cell cDNA amplification from tissue sections and provided an experimental platform conducive to high-quality transcriptomics, thereby revealing histology-associated transcriptomic heterogeneity in folliculogenesis in ovarian tissues.

## INTRODUCTION

Single-cell RNA sequencing (RNA-seq) has provided unprecedented opportunities for the study of cellular differentiations, states, and diseases in various biological fields, including developmental biology, stem cell biology, and reproductive medicine, since it was first achieved by using a quantitative cDNA amplification method and applied to mouse oocytes (Kurimoto et al. 2006; Tang et al. 2009). While many high-throughput single-cell RNA-seq methods have been developed (see (Svensson et al. 2018) for review), they typically lose histological information during cell dissociation from tissues. To preserve the histological information in transcriptome analyses, various *in situ* spatial transcriptomic methods that cover whole tissue sections have been developed (see (Liao et al. 2021) for review). These single-cell and spatial transcriptomics are extremely high throughput, rely on unique molecular identifiers or hybridization probes, and output relatively low transcriptomic contents with a low signal-to-noise ratio, in comparison with conventional deep RNA sequencing; usually, for example, the detectable number of genes ranges from hundreds to a few thousand, and exon-exon junctions and sequence variants are not identified (Waylen et al. 2020; Liao et al. 2021). As a result, the developed transcriptomics are optimal for identifying cell types in a large cell population and/or spatially annotating them, but are likely suboptimal for in-depth analyses of individual cells in tissues. For example, oogenesis undergoes quality control of oocytes during folliculogenesis accompanied by intimate interactions between oocytes and surrounding granulosa cells, and thus, understanding this process would require high quality single-cell transcriptomics tightly linked with histology (Zhang et al. 2018).

On the other hand, comprehensive, un-biased transcriptomics have been achieved and widely applied for arbitrarily targeted regions of interest (ROIs) isolated from tissue sections with laser capture microdissection (LCM) (Espina et al. 2006). LCM-based transcriptomics targets *in situ* cells/regions for deep RNA sequencing, but it has conventionally been used for bulk tissue fragments. On the other hand, several methods have been developed for LCM-based single-cell/low-input RNA-seq (Nichterwitz et al. 2016; Chen et al. 2017; Foley et al. 2019; Perez et al. 2021). Thus, this approach has advantages for use in the performance of unbiased, comprehensive transcriptomics in histologically identifiable small numbers of cells, including for the detection of exon junctions, and would be expected to provide information complementary to that of the currently available high-throughput single-cell RNA-seq and spatial transcriptomics (Liao et al. 2021). In addition, in situations calling for the analysis of cells/regions of interest across many sections, the targeting isolation strategy with LCM would be particularly cost effective.

In the earliest version of LCM-based single-cell RNA-seq, the cDNA amplification method for Smart-seq2 (Picelli et al. 2013) was directly applied to alcohol-fixed sections, with cell lysis using a non-denaturing detergent (Triton X-100) (Nichterwitz et al. 2016). Non-denaturing detergents are most frequently used for the lysis of freshly dissociated single cells, and such detergents can also be used in the subsequent enzymatic reactions for cDNA amplification in the same sampling tubes, a critical attribute for the success of low-input analyses such as analyses of single cells. On the other hand, the lysis efficiency of cells from sections with non-denaturing detergents has been controversial, which led Chen et al. (Chen et al. 2017) to propose a strategy that subjects a small number of cells in tissue sections to complete cell lysis under a denaturing condition followed by RNA purification with ethanol precipitation. Although efficient, high-quality RNA recovery is critical for quantitative transcriptome analysis of single cells, the laborious procedures involved in the RNA purification might limit the practical usability of such an approach (Le et al. 2015; Ghimire et al. 2021).

Moreover, formalin-fixed sections remain a challenge for high-quality, comprehensive single-cell RNA-seq, although formalin fixation achieves good tissue preservation and is widely used in histology (Titford and Horenstein 2005; Paavilainen et al. 2010). While a two-way RNA-seq method was developed for both alcohol- and formalin-fixed sections (Foley et al. 2019), in principal, it relied on an additional RNA hydrolysis for cell lysis and a short elongation time for cDNA amplification, thereby excluding transcript information other than the 3’-ends, and potentially compromising the sensitivity as well. Similarly, current single-cell transcriptomics often relies on the detection of 3’-ends or targeting probes, and frequently neglects additional sequence information such as exon–exon junctions.

Thus, an efficient, versatile cDNA amplification method for alcohol- and formalin-fixed tissue sections without RNA purification would enable comprehensive and robust *in situ* single-cell transcriptomics for histologically targeted cells of interest in a less labor-intensive manner. In this study, we developed cDNA amplification methods combined with an efficient cell lysis strategy for tissue sections, which uses a denaturing detergent for lysis, followed by quenching of the denaturing effect with an excess amount of a non-denaturing detergent (direct RNA recovery and quenching for LCM: DRaqL). The versatility of DRaqL was demonstrated by using it in combination with three different cDNA amplification protocols: SC3-seq (Kurimoto et al. 2006; Nakamura et al. 2015), Smart-seq2 (Picelli et al. 2013), and the protocol in the SMART-Seq v4 3’DE kit, which is a commercially available Smart-seq2-based kit that is compatible with multiplex cDNA library preparation and allows improved throughput (Takara Bio). The DRaqL-combined methods allowed efficient transcriptome profiling and exon–exon junction analyses of single cells isolated from alcohol-fixed sections with LCM comparable with those of freshly dissociated single cells. Furthermore, when combined with protease treatment, DRaqL was successfully applied to tissue sections strongly fixed with formalin (10%, 24 h at room temperature), enabling reliable single-cell RNA-seq from formalin-fixed sections.

By applying this method to mouse ovarian sections, we revealed a transcriptomic continuum of growing oocytes and detected splice isoforms important in oogenesis. We constructed a statistical model of the transcriptome of oocytes based on their size, and, by examining deviations from the model, we found heterogeneity of the size–transcriptome relationship in oocytes. Moreover, we revealed genes that were differentially expressed in granulosa cells in association with their histological parameters. We thus developed a versatile, effective single-cell cDNA amplification strategy for high-quality RNA-seq from alcohol- and formalin-fixed tissue sections, and revealed histology-associated transcriptomic heterogeneity in folliculogenesis in mouse ovarian tissues.

## RESULTS

### Experimental system for quantitative examination of cDNA amplification from sections

First, we sought a method to efficiently amplify single-cell cDNAs from alcohol-fixed sections. To circumvent cellular heterogeneity in tissues, we set up a system to evaluate the cDNA amplification of single cells isolated with LCM from the alcohol-fixed, stained sections of frozen-cell blocks composed of homogeneous cultured cells—namely, embryonic stem cells under a 2i-LIF condition (2i-LIF mESCs) (Ying et al. 2008; Marks et al. 2012) (Fig. 1A, B). As a gold standard, fresh single cells from the same culture batches were also dissociated and isolated. We amplified cDNAs of these single cells by means of the amplification protocol used in the SC3-seq method (Kurimoto et al. 2006; Kurimoto et al. 2008; Nakamura et al. 2015), and compared their gene expression levels using real-time PCR.

**Figure 1.**
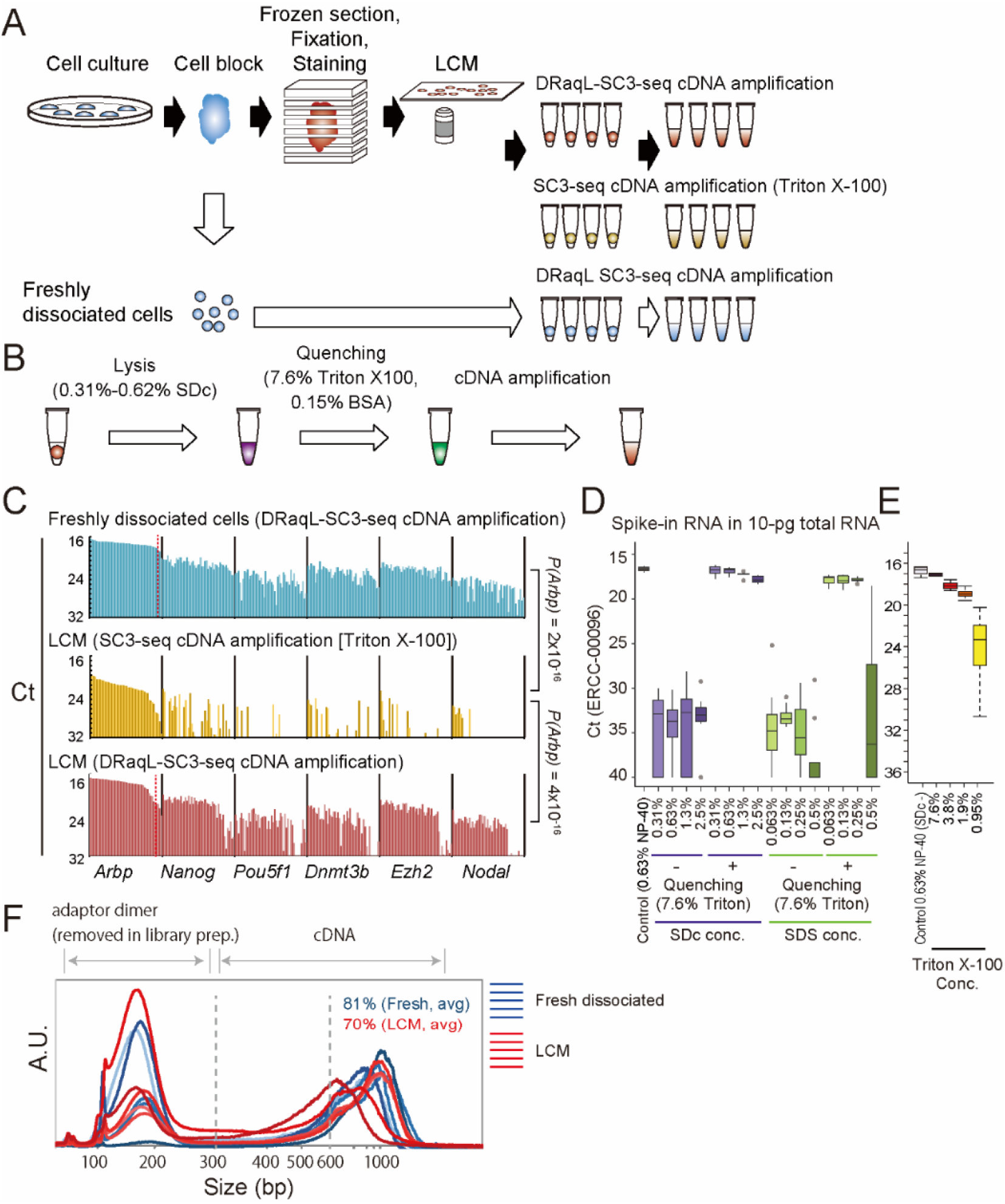
Single-cell DRaqL-SC3-seq cDNA amplification from cell blocks. (A) Schematic representation of the development of the DRaqL-SC3-seq cDNA amplification method. (B) Schematic representation of DRaqL-combined cDNA amplification. (C) Real-time PCR analysis of cDNAs of freshly dissociated single mESCs (top), single cells isolated from alcohol-fixed sections of cell blocks, lysed using Triton X-100 (middle) and DRaqL (bottom) (n = 48 each). cDNAs were amplified with the indicated methods. *P* values by a *t*-test for *Arbp C*_t_ values are also shown. (D) Evaluation of the SC3-seq cDNA amplification efficiency from 10-pg RNA of mESCs with different concentrations of SDc and SDS, with and without quenching by 7.6% Triton X-100. *C*t values of spike-in ERCC-00096 RNA (∼3,000 copies) are represented with boxplots (n = 8 each). (E) Evaluation of cDNA amplification efficiency with different concentrations of Triton X-100 for quenching of 0.63% SDc (n = 4 each). (F) Electropherograms of cDNAs analyzed using Bioanalyzer (Agilent Technologies). The adaptor dimers (<300 bps) are removed during library preparation.

We first evaluated embedding media, and found that OCT compound, a widely used embedding medium, compromised the gene expression profiles compared with 10% PVA (Supplemental Fig. S1A, B). Thus, we decided to use 10% PVA for embedding.

### Efficient cDNA amplification with DRaqL-SC3-seq

Next, we evaluated the cDNA amplification using a non-denaturing detergent, Triton X-100 (0.63%; SC3-seq cDNA amplification [Triton X-100]), for single cells isolated from sections of the cell blocks. In agreement with the previous study (Chen et al. 2017), the cells isolated from the cell-block sections showed significantly reduced expression levels of *Arbp*, a highly expressed housekeeping gene, compared with the freshly dissociated cells (Fig. 1C).

We therefore hypothesized that the use of denaturing detergents would improve the cell lysis efficiency, and that subsequent enzymatic reactions would be allowed by quenching the denaturing effect with the addition of an excess amount of non-denaturing detergents (Fig. 1B). We evaluated the quenching effect with cDNA amplification from the single-cell equivalent amount (10 pg) of total RNA, and found that the denaturing detergents sodium deoxycholate (SDc) (≤0.63%) and sodium dodecyl sulfate (SDS) (≤0.25%) were efficiently quenched by the addition of 7.6% Triton X-100 (Fig. 1D, E). In addition, efficient quenching required bovine serum albumin (BSA) at a concentration up to 0.15% (Supplemental Fig. S1C). Thus, we decided to use 0.31%–0.63% SDc for the cell lysis, and 7.6% Triton X-100 and 0.15% BSA for the quenching, and we termed the lysis method “<u>D</u>irect <u>R</u>NA recovery <u>a</u>nd <u>q</u>uenching for <u>L</u>CM” (DRaqL). When we combined DRaqL with the SC3-seq cDNA amplification, we used 0.63% SDc for cell lysis.

Next, we evaluated the DRaqL-SC3-seq cDNA amplification using single cells isolated from the alcohol-fixed sections of the cell blocks (48 cells each). As shown in Figure 1C and Supplemental Figure S1D, the amplification efficiency was significantly improved compared with the efficiency of SC3-seq cDNA amplification (Triton X-100) (*P* = 4×10^−16^), and was similar to the amplification efficiency from freshly dissociated cells, albeit with a ∼19% reduction in the success rate (Supplemental Fig. S1E).

We also found that the electropherograms of amplified cDNAs were similar between these cells, with only a small size reduction in the cell-block cells (∼81% and ∼70% of cDNA >600 bps, respectively) (Fig. 1F). Thus, the DRaq-SC3-seq cDNA amplification is a useful single-cell cDNA amplification method from alcohol-fixed sections.

### Examination of transcriptome profiling with DRaqL-SC3-seq from alcohol-fixed sections

Next, we examined the amplified cDNA in a genome-wide manner, using the 3’-sequencing method, SC3-seq (Nakamura et al. 2015; Nakamura et al. 2016) (Fig. 2A). We found that the freshly dissociated cells and cell-block cells showed similar profiles of the number of detected genes (10,730 and 9,886 genes on average, respectively) (Fig. 2B, C). The expression levels of spike-in RNAs (ERCCs) were also essentially the same in both cells (Supplemental Fig. S2A, B). Scatterplots of gene expression levels in single cells showed no large difference between these types of cells (Fig. 2D). Consistent with the above findings, principal component analysis (PCA) showed that about 40% of the 97%-confidence interval ellipse areas were overlapped between the freshly dissociated cells and cell-block cells (Fig. 2E). In addition, the average gene expression patterns were similar between the freshly dissociated cells and cell-block cells, with only small systematic errors, as described below (Fig. 2F). These results demonstrate that DRaqL-SC3-seq allows a high-quality single-cell transcriptome analysis for alcohol-fixed sections, albeit accompanied by non-negligible artifacts.

**Figure 2.**
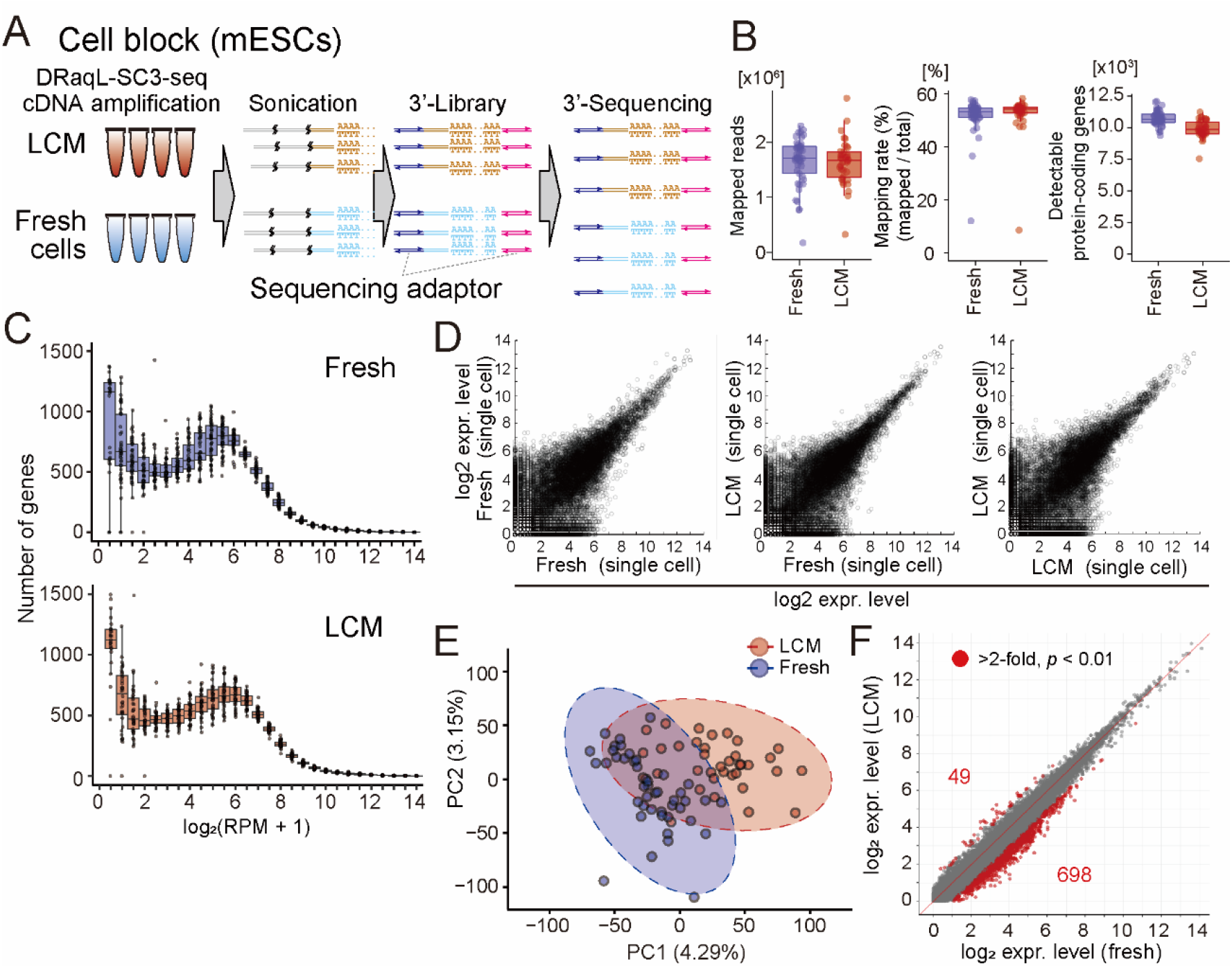
Single-cell transcriptome analysis with DRaqL-SC3-seq. (A) Schematic representation of the 3’-sequencig with DRaqL-SC3-seq. (B) Box plots showing the numbers of mapped reads (left), mapping rates (middle), and detectable protein-coding genes (>0 mapped reads) (right). Freshly dissociated single cells (Fresh) and single cells from alcohol-fixed cell-block sections are shown (LCM). (C) Frequency plots of gene expression levels in freshly dissociated single cells (Fresh) (top), and single cells from alcohol-fixed cell-block sections (LCM) (bottom). (D) Representative scatterplots of gene expression levels between freshly dissociated single cells (Fresh) and single cells isolated from alcohol-fixed cell-block sections (LCM). (E) PCA of freshly dissociated single cells (Fresh) and single cells isolated from alcohol-fixed cell blocks (LCM). 97%-confidence interval ellipses were represented with dashed lines. (F) Scatterplots of the averaged log_2_ (RPM+1) values between freshly dissociated single cells and single cells isolated from alcohol-fixed cell-block sections. Differentially expressed genes (log_2_ difference >1, and *p*<0.01 by *t*-test) are indicated with red circles.

### Evaluation of errors and biases caused by the use of alcohol-fixed sections

To dissect the errors and biases in DRaqL-SC-3seq from the alcohol-fixed sections, we performed an in-depth comparison of transcriptome between the freshly dissociated and cell-block cells, by taking advantage of the fact that these cells were prepared from the same culture batch of homogeneous 2i-LIF mESCs.

We examined the statistical significance of differences in the detection rate of each gene (i.e., the frequency of cells in which expression of the gene was detected), between these two types of cells. We found that, out of the genes detected in at least one sample (18,992), 5% showed a significant reduction of detection rates in the cell-block cells (*p<*0.01; detection-rate errors) (Supplemental Fig. S3A, B). The vast majority of the detection-rate errors occurred in genes with relatively low expression levels (95% occurred in log_2_ [reads per million mapped reads (RPM) +1] <4; i.e., <15 copies/cell).

Next, as mentioned above, we examined the differences of average expression levels between these cells, and found that 0.26% and 3.7% of genes were up- and down-regulated, respectively, in the cell-block cells by >2 fold with *p<*0.01 (expression-level biases) (Fig. 2F). These biases were distributed in relatively low expression levels (54% and 97% occurred in log_2_ [RPM+1] <4 and <6, respectively) (Supplemental Fig. S3C, D). Therefore, DRaqL-SC3-seq showed both detection-rate errors and expression-level biases due to the use of alcohol-fixed sections in only a small fraction of lowly expressed genes overall.

### Exon–exon junction analysis of DRaqL-SC3-seq cDNA amplification

We next asked whether DRaqL SC3-seq cDNA amplification from the sections was applicable to quantitative expression profiling of exon–exon junctions, by applying the whole cDNAs to sequencing with the Y-adaptor sequencing (Fig. 3A; Supplemental Fig. S4).

**Figure 3.**
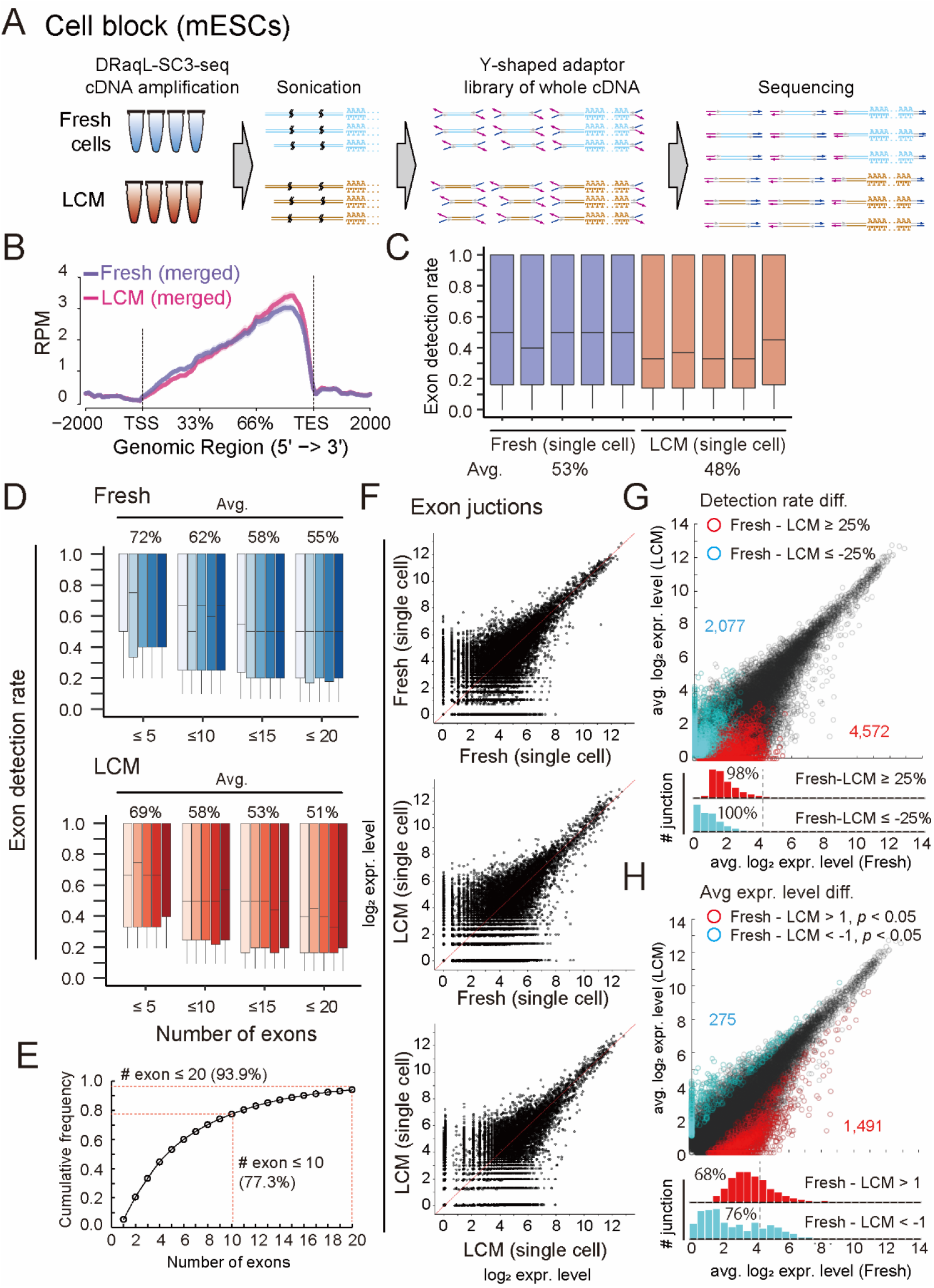
Exon–exon junction profiling of the DRaqL-SC3-seq cDNA amplification. (A) Schematic representation of the Y-shaped adaptor sequencing of the amplified cDNAs. (B) Numbers of mapped reads plotted between the transcription start site (TSS) and transcription end site (TES). The averages for freshly dissociated single cells (Fresh) and single cells from alcohol-fixed cell-block section (LCM) are shown (n = 5 each). (C) Box plots of detection rates of exons in single cells (n = 5 each). (D) Box plots of exon detection rates of genes with different numbers of exons (indicated number of exons or fewer) in single cells (n = 5 each). (E) Frequency of mouse protein-coding genes that have the indicated numbers of exons or fewer. (F) Representative scatterplots of the expression levels of exon–exon junctions between freshly dissociated single cells (Fresh) and single cells from alcohol-fixed cell-block sections (LCM). (G, H) Scatterplots of averaged exon–exon junction expression levels of freshly dissociated single cells (Fresh) and single cells from alcohol-fixed cell-block sections (LCM) (n = 5 each). Exon–exon junctions for which the detection-rate difference was ≥25% (G) and the log_2_ expression difference >1 (*p<*0.05) (H) are indicated with red and cyan. Histograms of these exon–exon junctions are shown below the scatterplots.

The mapping profiles of the Y-adaptor sequencing for these cDNAs showed a bias toward the 3’-ends, while even the near 5’-end regions showed mapped reads (Fig. 3B). To calculate the exon detection rates in expressed genes, we counted the number of detectable exons for each sample (Trimmed Mean of M-values [TMM] >2). In the freshly dissociated cells, we detected an average of 53% of exons (Fig. 3C). For protein-coding genes containing ≤20 exons, 55% exons were detected (Fig. 3D). It is worth noting that the numbers of exons were ≤20 in ∼94% of mouse genes. Thus, the SC3-seq cDNA amplification method successfully detected about half of the exons in most genes in the freshly dissociated single cells.

Then, we examined the profiles of exon detection rates in the cell-block cells, and found a 5% reduction of exon detection rates (48%) compared with the freshly dissociated cells (Fig. 3C). In genes with ≤20 exons, more than half of exons were detected (51%) (Fig. 3D). This indicates that the Y-adaptor sequencing allows exon profiling in alcohol-fixed sections at an efficiency comparable with that in freshly dissociated cells.

Next, we counted the reads mapped to the exon–exon junctions, and quantified their expression levels (Fig. 3F). In total, we detected 25,283 junctions expressed in at least one sample of freshly dissociated and cell-block cells. Scatterplots showed that the junctional expression levels in single cells were similar between these types of cells. Next, we examined differences in the detection rates and expression levels between freshly dissociated and cell-block cells. We found that vast majority of >25% difference in detection rates of junctions occurred at a log_2_ expression level <4 (Fig. 3G). Similarly, differences in expression levels (>2-fold, *p<*0.05) were mainly observed in junctions with a log_2_ expression level <4 (Fig. 3H). Thus, these errors and biases in the exon– exon junction profiling showed trends similar to those in the 3’-sequencing.

Collectively, these results indicate that the DRaqL-SC3-seq cDNA amplification is compatible with a high-quality single-cell transcriptome analysis and exon–exon junction profiling from the alcohol-fixed sections, albeit with bias and errors in the low range of expression levels.

### Application of DRaqL-SC3-seq to mouse ovarian sections

Next, we examined whether DRaqL-SC3-seq can address biological questions, by applying it to alcohol-fixed sections of mouse ovaries (Fig. 4A). We analyzed the transcriptome of 44 single growing oocytes from primary-to-early antral follicles and 57 single granulosa cells in secondary-to-early antral follicles, isolated with LCM from the sections (Supplemental Fig. S5A−C). The numbers of detectable genes in single oocytes and granulosa cells were comparable with those in a previous study of the single-cell transcriptome of freshly dissociated, human oocytes and granulosa cells, respectively, analyzed with Smart-seq2 (Fan et al. 2021). This suggests that DRaqL-SC3-seq showed sufficient sensitivity for single cells in tissue sections (Supplemental Fig. S5B, D). PCA showed that these cell types formed clearly distinct clusters, wherein PC1 represented the difference between oocytes and granulosa cells (Fig. 4B). In addition, scatterplots between serial sections from the same oocytes demonstrated reproducibility of the analyses (Supplemental Fig. S6).

**Figure 4.**
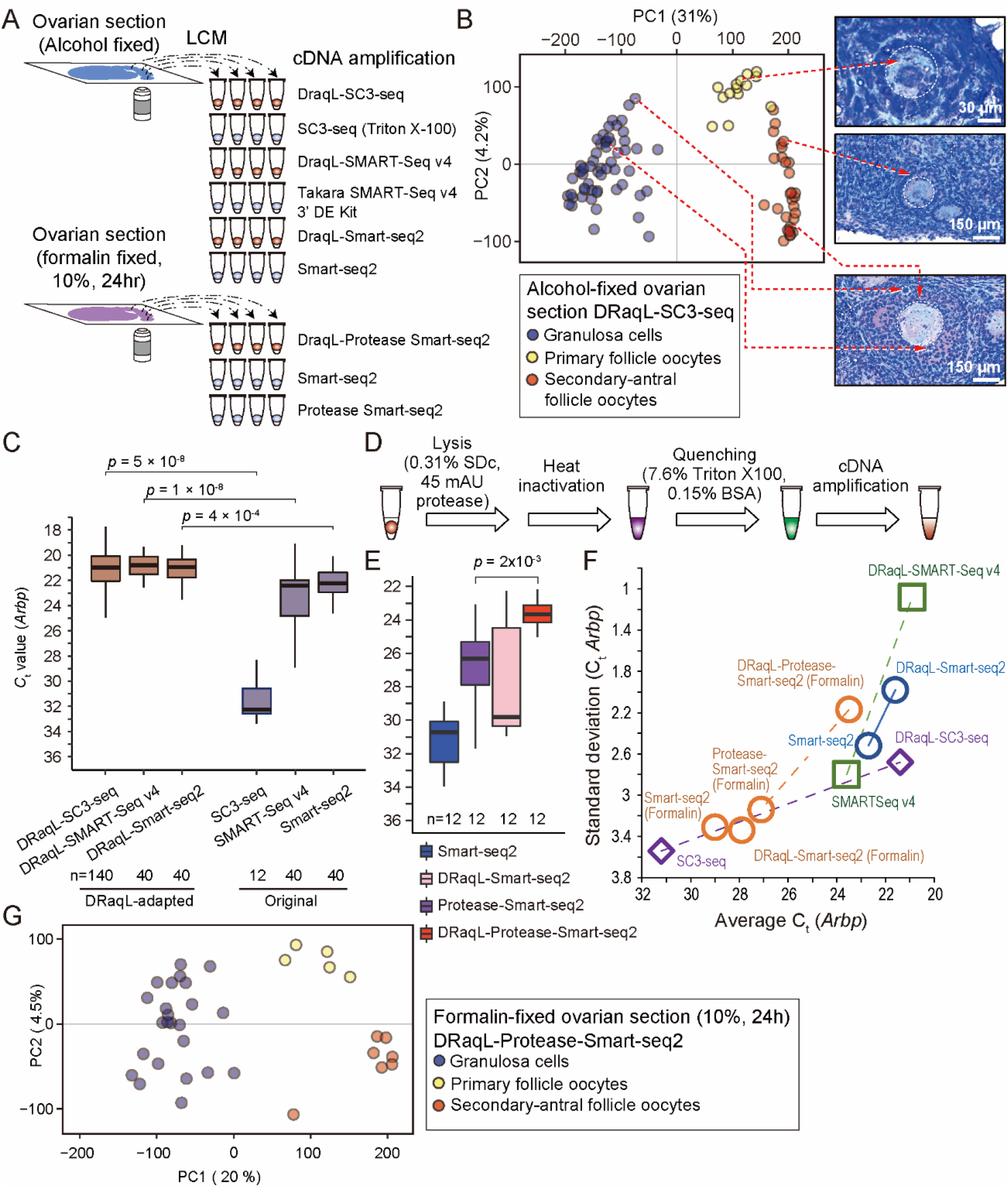
Application of DRaqL-adapted cDNA amplification methods to mouse ovarian sections and their expansion to formalin-fixed sections. (A) Schematic representation of different cDNA amplification methods for single cells isolated from alcohol- and formalin-fixed ovarian sections. (B) PCA of single oocytes and granulosa cells isolated from mouse ovarian sections analyzed with DRaqL-SC3-seq. Representative histological images are shown to the right of the plots, and isolated cells are indicated with dashed lines. (C) Real-time PCR analysis of cDNAs amplified from single granulosa cells isolated from alcohol-fixed ovarian sections. *Ct* values of *Arbp* in cDNAs amplified with indicated methods are represented with boxplots. *P* values by a *t*-test are also shown. (D) Schematic representation of DRaqL combined with protease treatment. (E) Real-time PCR analysis of cDNAs amplified with different Smart-seq2-based cDNA amplification methods from single granulosa cells isolated from formalin-fixed ovarian sections. The *Arbp C*_t_ values are represented with boxplots. *P* values by a *t*-test are shown above the graph. (F) Scatterplots of the averages and standard deviations of the *Arbp C*_t_ values in cDNAs amplified with the indicated methods from alcohol- and formalin-fixed sections. The DRaqL-adapted methods are linked with their original methods with dashed lines. The average and standard deviation for DRaqL-Protease-Smart-seq2 and Smart-seq2 from formalin-fixed sections were calculated for the three independent experiments shown in Supplemental Figure S8F. (G) PCA of single oocytes and granulosa cells isolated from the formalin-fixed mouse ovarian sections. The color coding is the same as in (B).

We also performed RNA-seq of pooled granulosa cells isolated from the sections (10, 100 – 300, and >300 cells) using the DRaqL-SC3-seq protocol with reduced numbers of PCR cycles, and found that the number of detectable genes increased according to the number of pooled cells (Supplemental Fig. S7A, B). Thus, we concluded that DRaqL-SC3-seq is applicable to quantitative transcriptome analysis for single cells and arbitrary sizes of ROIs isolated from alcohol-fixed, mouse ovarian sections.

### Application of DRaqL to other downstream cDNA amplifications

Next, for single granulosa cells isolated from the ovarian sections, we asked whether other cDNA amplification methods can be combined with DRaqL (Fig. 4A). First, we examined the use of DRaqL in conjunction with a SMART-Seq v4 3’DE Kit (Takara Bio), a commercially available Smart-seq2-based kit that allows multiplex cDNA library preparation by including index sequences in the reverse transcription primers. We found that its cDNA amplification efficiency was reduced with the use of DRaqL (Supplemental Fig. S8A). In addition, the efficiency was slightly better when the SDc concentration was 0.31% than when it was 0.63%. Thus, using the cell lysis buffer containing 0.31% SDc, we examined the use of additional reverse transcriptases with this kit, and found that the optimal combination (SuperScript II and SuperScript III) is capable of cDNA amplification with efficiency and stability similar to those of the original protocol (Supplemental Fig. S8B). The cDNA amplification efficiency for single granulosa cells isolated from the alcohol-fixed ovarian sections was similar to that of DRaqL-SC3seq, and we termed this method DRaqL-SMART-Seq v4 cDNA amplification (Fig. 4C).

Next, using the combination of separately available reagents and oligo nucleotides with multiplex index sequences, we developed another Smart-seq2-based cDNA amplification method adapted to DRaqL (DRaqL-Smart-seq2), and found that this combination achieved cDNA amplification efficiency similar to those by DRaqL-SC3-seq and DRaqL-SMART-Seq v4 (Fig. 4C). In fact, for single granulosa cells in the alcohol-fixed sections, DRaqL-SMART-seq v4 and DRaqL-Smart-seq2 improved both the efficiency and stability of cDNA amplification over SMART-Seq v4 3’DE Kit, Smart-seq2, and other LCM-combined cDNA amplification methods (Nichterwitz et al. 2016; Chen et al. 2017; Foley et al. 2019) (Fig. 4C, F; Supplemental Fig. S8C, D). Thus, we established DRaqL-adapted, efficient single-cell Smart-seq2-based methods for alcohol-fixed tissue sections.

### Application of DRaqL to formalin-fixed sections

Next, we examined whether DRaqL-adapted cDNA amplification is compatible with formalin-fixed sections. To test the versatility of the method, we used a strong fixative condition, 10% formalin for 24 h at room temperature, which is frequently used in histopathology (Fig. 4A, D). We applied DRaqL-Smart-seq2 and Smart-seq2 to mouse ovarian sections fixed with this condition, and found a severe reduction of cDNA amplification efficiency (Fig. 4E).

To digest the formalin-fixed cellular components, we thus combined DRaqL-Smart-seq2 with a thermolabile protease with an optimization of heat inactivation (DRaqL-Protease-Smart-seq2) (Fig. 4E; Supplemental Fig. S8E). Application of DRaqL-Protease-Smart-seq2 to single granulosa cells from the formalin-fixed sections showed robustly improved cDNA amplification efficiency over those of Smart-seq2, DRaqL-Smart-seq2, and another previous method exploiting proteinase K and additional RNA hydrolysis for cell lysis (Foley et al. 2019) (Fig. 4E, F; Supplemental Fig. S8F).

To evaluate the effect of DRaqL on the protease treatment more directly, we compared the cDNA amplification efficiency of DRaqL-Protease-Smart-seq2 with a simple combination of protease and Smart-seq2 (Protease-Smart-seq2) (Perez et al. 2021). While the amplification efficiency of Protease-Smart-seq2 was better than those of Smart-seq2 and DRaqL-Smart-seq2, DRaqL-Protease-Smart-seq2 showed much better cDNA amplification efficiency and stability than Protease-Smart-seq2, demonstrating that DRaqL significantly improved the efficiency of protease treatment (Fig. 4E, F).

Importantly, DRaqL-Protease-Smart-seq2 showed reproducible cDNA amplification efficiency from formalin-fixed sections, with a standard deviation comparable to that of DRaqL-Smart-seq2 from alcohol-fixed sections (Fig. 4F). RNA-seq of single oocytes and granulosa cells from the formalin-fixed sections with DRaqL-Protease-Smart-seq2 showed gene expression profiles similar to those from alcohol-fixed sections with DRaqL-SC3-seq (Fig. 4G; Supplemental Fig. S8G–I). These data demonstrate that DRaqL-Protease-Smart-seq2 is capable of robust, high quality transcriptome analysis for single cells in sections strongly fixed with formalin.

### Morphology-associated transcriptome dynamics of oocytes in follicles

Next, we asked how the morphology of oocytes is associated with their transcriptome, using DRaqL-SC3-seq. Along the PC2 axis of the transcriptome, the growing oocytes showed expression profiles tightly linked with the follicular morphology, forming clearly different clusters between primary follicles and secondary-to-antral follicles (Fig. 4B). Genes previously known to be involved in oogenesis (such as *Obox, Oog, Oosp1, Bmp15, Gdf9, Izumo1r, H1foo*, and *Bcl2l10*) contributed highly to PC2, suggesting that it represented the growth axis of oocytes (Supplemental Table S4). More importantly, the PC2 values were highly correlated with the sizes of oocytes and follicles (rank correlation coefficient −0.83 for both), indicating a quantitative association between the histological parameters and transcriptome (Fig. 5A; Supplemental Fig. S9; Supplemental Table S7).

**Figure 5.**
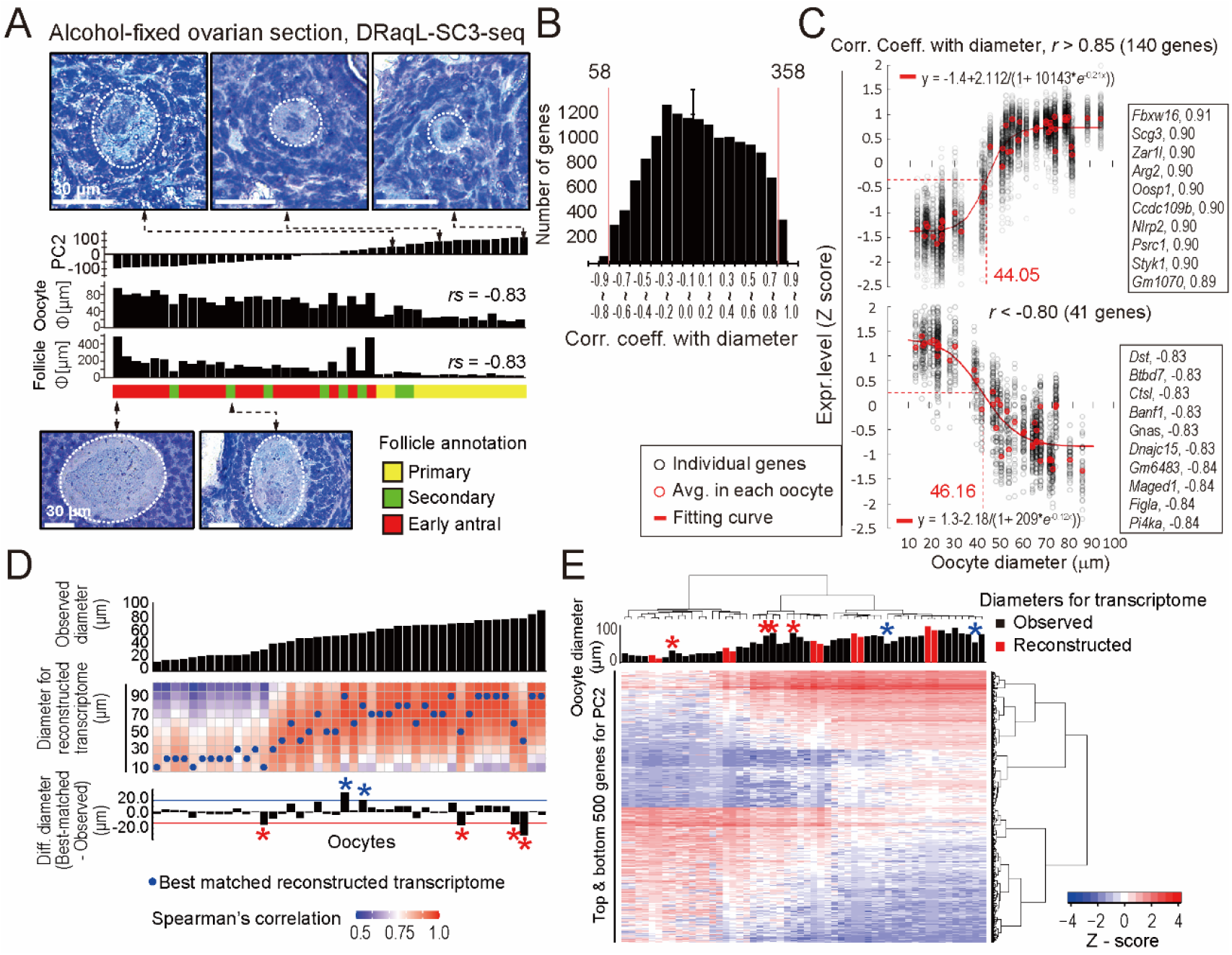
Morphology-associated transcriptome difference of oocytes revealed with DRaqL-SC3-seq. (A) Relationship of PC2 values of oocytes isolated from the alcohol-fixed sections and their morphology. PC2 values (top), oocyte diameters (middle), and follicle diameters (bottom) are shown in the bar graphs. The annotations of follicle stages are color coded. Representative histological images of follicles are also shown. The Spearman’s rank correlation efficient (*rs*) between PC2 and the size of oocytes and follicles is indicated. (B) Histogram of Pearson’s correlation coefficient between the expression levels of individual genes and oocyte diameters. (C) Scatterplots of log_2_ gene expression levels against oocyte diameters. Positively correlated genes (top, *r* >0.85, n = 140) and negatively correlated genes (bottom, *r* < -0.80, n = 41) are shown. Average *Z* scores of these genes in individual oocytes and their fitting curves are plotted with red open circles and red lines, respectively. Gene symbols and *r* values of the top 10 genes are also indicated. (D) Heat map representation of correlation coefficients between the transcriptome of individual oocytes and reconstructed transcriptome model for indicated diameters. Diameters of oocytes are shown with the bar graphs. Blue dots indicate the reconstructed transcriptomes best-matched with individual oocytes (i.e., the highest correlation). The difference of diameters between the individual oocytes and the best-matched models are shown with the bar graphs (differences >20 μm are indicated with red and blue asterisks). (E) Heat map representation of expression levels of the top 500 genes with positive and negative PC2 values. Diameters of oocytes are shown with the bar graphs, and those of reconstructed transcriptome model are indicated with red shading. Asterisks indicate the same oocytes as in (E).

Next, therefore, we investigated the correlations between the expression levels of individual genes and the oocyte diameters (Fig. 5B). The expression levels of the genes highly positively and negatively correlated with the diameters (*r*>0.85 and *r*<-0.80, respectively) were continuously distributed, and sigmoid curves fitting their distribution showed inflection points at similar diameters (44 μm and 46 μm, respectively) (Fig. 5C; Supplemental Table S5). This suggests that oocytes change their transcriptome most dramatically when their size has grown to around these values, reflecting primordial-to-primary follicle transition (Pan et al. 2005; Hamazaki et al. 2021).

Then, we evaluated the relationship between the transcriptome of oocytes and their diameter by constructing a statistical model (Fig. 5D, E; Supplemental Fig. S9A). Using simple regression analyses with PC1 and PC2 as objective variables and diameter as an explanatory variable, we reconstructed the transcriptome of oocytes for every 10 μm of diameter, and examined which reconstructed transcriptome data were best matched with individual oocytes (Fig. 5D). We found that the reconstructed transcriptome was best matched with the transcriptome of oocytes with the most similar diameters (Fig. 5D, E), suggesting that PC1 and PC2 had sufficient information to link gene expression and the size of oocytes. On the other hand, in 4 out of the 44 oocytes examined, we found that this statistical model yielded data widely discrepant from the observed data; the diameters of the best-matched, reconstructed transcriptome were significantly smaller than the diameters of the observed oocytes (>20 μm decrease; Fig. 5D). In line with these observations, these oocytes showed expression signatures similar to those of smaller oocytes (Fig. 5E). These results indicate that our model successfully reconstructed the transcriptome in the development of normal oocytes, and the deviations from the model delineate heterogeneity of their size–transcriptome relationship, likely reflecting the underlying selective processes for dominant follicles (Deane 1952; Byskov 1974).

### Detection of splicing isoforms in single oocytes in sections

Next, we conducted an exon–exon junction analysis of the oocytes in the alcohol-fixed sections, because there is growing evidence of the importance of alternative splicing in oocytes for meiotic progression, oocyte growth and maturation, and female fertility (Tang et al. 2009; Do et al. 2018; Kasowitz et al. 2018; Cheng et al. 2020; Li et al. 2020; Yu et al. 2021) (Supplemental Fig. S10). We applied the Y-adaptor sequencing to the single-oocyte cDNAs (n = 5) (Supplemental Fig. S9C), and successfully detected splice isoforms in many genes, including *H1foo, Lsm14b, Rac1, Trp53bp1, Oosp1*, and *Parl*. For example, the minor splice isoforms of *Lsm14b, Rac1*, and *Trb53bp1*, which are regulated by ESRP1 in oocytes (Yu et al. 2021), were detected in this analysis. Moreover, in *Serf2* and *Cox6b2*, we detected oocyte-specific minor splice isoforms.

### Histology-associated gene expression differences in granulosa cells

Finally, we investigated relationships between the histology and transcriptome of granulosa cells in early antral follicles. Granulosa cells have direct contacts with oocytes through gap junctions on transzonal projections, depending on their positions relative to oocytes (Simon et al. 1997; Li and Albertini 2013). Thus, we asked whether or not the gene expression profiles of granulosa cells were different between the cells that histologically neighbored oocytes and those that did not— i.e., whether there were differentially expressed genes (DEGs) between neighboring (n = 31) and non-neighboring granulosa cells (n = 26) (Fig. 6A). The neighboring granulosa cells formed a layer covering the surface of the oocyte, while the non-neighboring granulosa cells formed relatively loosened structures compatible with the formation of follicular cavities (Supplemental Fig. S5A, S9B).

**Figure 6.**
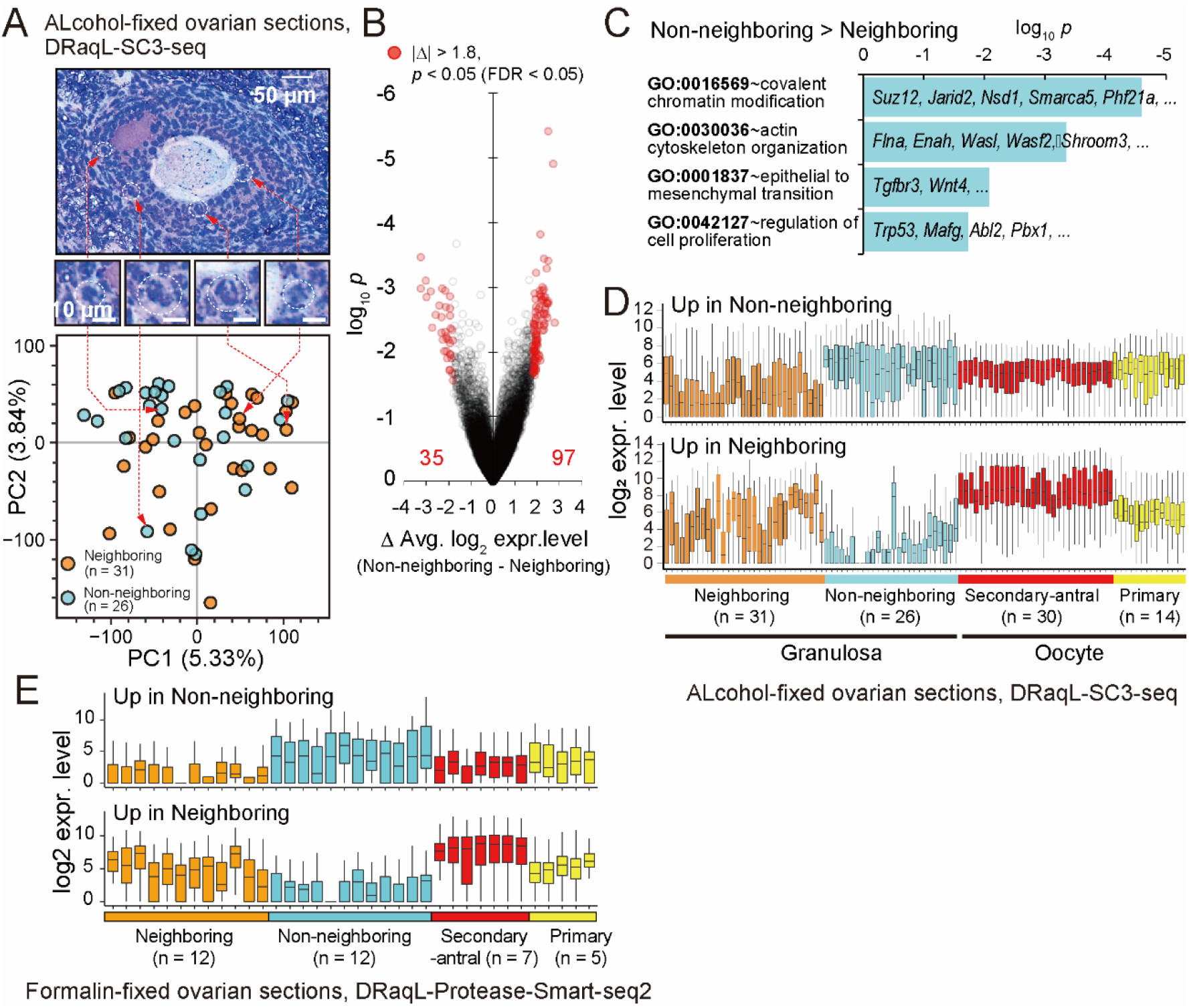
Single-cell transcriptome analysis of granulosa cells in ovarian sections with DRaqL-SC3-seq. (A) PCA of DRaqL-SC3-seq transcriptome data of granulosa cells in alcohol-fixed sections. Cells neighboring oocytes (Neighboring) and those not neighboring oocytes (Non-neighboring) are indicated with orange and cyan circles, respectively. (B) Volcano plots of log_2_ expression level differences and log_10_ *p* values between neighboring and non-neighboring granulosa cells. Genes with differences in log_2_ expression levels of more than 1.8 (i.e., >3.5 fold) and *p<*0.05 with FDR <0.05 are indicated with red circles. (C) Bar graphs showing enrichment of gene ontology (GO) terms in genes up-regulated in non-neighboring granulosa cells. (D) Box plots of the log_2_ expression levels of genes up-regulated in the neighboring and non-neighboring granulosa cells. Expression levels in single oocytes and granulosa cells are shown. (E) Box plots of the log_2_ expression levels of the genes shown in (D) in single oocytes and granulosa cells isolated from the formalin-fixed sections (analyzed with DRaqL-Protease-Smart-seq2).

As shown in Figure 6A, these two types of cells showed no clear transcriptome differences in PCA. However, a direct comparison of gene expression levels revealed 35 and 97 genes that were up-regulated in neighboring and non-neighboring granulosa cells, respectively (average log_2_ difference >1.8 [i.e., >∼3.5-fold], FDR<0.05) (Fig. 6B; Supplemental Table S6). We found that genes up-regulated in non-neighboring granulosa cells were enriched with genes coding for chromatin modifiers, including Polycomb Repressive Complexes (*Suz12, Jarid2*), regulators of chromatin modifiers (*Ogt, Nsd1*), a histone H3-lysine4-mediated regulator (*Ph421a*), and a SWI/SNF family member (*Smarca5*) (Fig. 6C). In addition, these cells were also enriched with genes involved in actin cytoskeleton organization (*Flna, Enah*), epithelial-to-mesenchymal transition (*Tgfbr3, Wnt4*), and regulation of cell proliferation (*Trp53, Abl2*), suggesting that their histological positions were associated with distinct proliferative and morphological characters, indicative of granulosa cell differentiation.

On the other hand, the 35 genes up-regulated in the neighboring granulosa cells showed highly heterogeneous expression levels even among these cells, while they were nearly undetectable in the non-neighboring granulosa cells (Fig. 6D). Interestingly, they included oocyte-specific genes, such as *Obox* family members, which were expressed in oocytes in the secondary-to-early antral follicles at much higher expression levels. These results most likely reflected gap junction communications in the oocyte and granulosa cells (Anderson and Albertini 1976; Simon et al. 1997; Matzuk et al. 2002).

RNA-seq of 10 pooled granulosa cells from early antral follicles confirmed the expression profiles of these DEGs, supporting the accuracy of the histology-linked gene expression analysis at the single-cell level with DRaqL-SC3-seq (Supplemental Fig. S7A, C). In bulk RNA-seq of whole granulosa from early antral follicles, these DEGs were expressed at intermediate levels, consistent with the idea that bulk granulosa is a mixture of these cell types. These results demonstrate that these DEGs can be identified only by identifying their histological affiliation to oocytes followed by a transcriptome analysis.

Additionally, DRaqL-Protease-Smart-seq2 showed expression profiles of these DEGs in the single granulosa cells from the formalin-fixed sections similar to those in the single granulosa cells from the alcohol-fixed sections, again demonstrating the robust performance of this method for the formalin-fixed sections (Fig. 6E).

We further examined the expression profiles of these DEGs in previously published single-cell transcriptome data for mouse ovarian somatic cells (Li et al. 2021) and human cumulus oocyte complexes (Fan et al. 2021). We found antagonistic expression patterns of these DEGs in subsets of granulosa cells in the public datasets (Supplemental Fig. S11). These results demonstrate that DRaqL-SC3-seq for ovarian sections revealed previously unidentified co-expression patterns in granulosa cells.

Collectively, these results demonstrated that single-cell RNA-seq with DRaqL from ovarian sections allowed quantitative analyses of histology-based expression differences that cannot be revealed by transcriptomics alone, as well as splice isoform analyses.

## DISCUSSION

In this study, we developed robust quantitative cDNA amplification methods combined with cell lysis by DRaqL for single cells isolated from alcohol- and formalin-fixed sections. The results of quantitative evaluation revealed that DRaqL-SC3-seq from alcohol-fixed sections showed only small errors and biases occurring in relatively lowly expressed genes, and showed amplification efficiency similar to those of the DRaqL-adapted Smart-seq2-based methods, suggesting the versatility of DRaqL.

Combining DRaqL with protease treatment also enabled robust cDNA amplification and RNA-seq from sections fixed with formalin, the most popular fixative for tissue preservation (Titford and Horenstein 2005; Paavilainen et al. 2010) (Fig. 4). The strong fixative condition used in this study, 10% formalin for 24 h at room temperature, is preferred in histopathology (Titford and Horenstein 2005). Both cDNA amplification efficiency and stability with our method were significantly improved compared with the previous approaches, suggesting that DRaqL would facilitate cell lysis by protease.

For efficient cell lysis compatible with subsequent enzymatic reactions, DRaqL takes advantage of the efficient incorporation of a small amount of denaturing detergent into micelles of non-denaturing detergents (Jonstromer and Strey 1992; Sivars et al. 1994). This strategy has been used for the extraction of bacterial genomic DNA with SDS, followed by region-specific PCR after quenching with Tween20 (Goldenberger et al. 1995). This tactic has also been employed in the Hi-C technique, in which SDS is used to de-condense genomic DNAs followed by quenching with Triton X-100, in preparation for subsequent reactions such as restriction enzyme treatment (Lieberman-Aiden et al. 2009). As another example, a mix of SDS and non-ionic detergents has been used for cellulase-mediated glycolysis (Eriksson et al. 2002). In this study, we applied the combination of a denaturing detergent and quenching to RNA-seq of alcohol- and formalin-fixed sections and rigorously evaluated the performance of the method, expanding the application range of this principle in quantitative biology.

By applying this method to alcohol-fixed sections of mouse ovaries, we successfully depicted histology-associated transcriptional differences of oocytes and granulosa cells (Figs. 4–6). The formation of discrete clusters of oocytes in primary follicles and secondary-to-early antral follicles may reflect transcriptomic changes during the primary-to-secondary follicle transition (Williams and Erickson 2000; Zhang et al. 2018). The quantitative continuum of the size– transcriptome relationships of oocytes allows us to realize morphology-based prediction of the transcriptome (Fig. 5), providing a significantly higher resolution of this relationship than in a previous study in which dissociated oocytes were classified by diameter into three clusters and the differentially expressed genes were identified (Gu et al. 2019). It would be worth noting that the transcriptome data obtained in this study were associated with the intact morphology of snap-frozen ovarian tissues, without any artifacts caused by cell dissociation. The heterogeneity of the relationship between the transcriptome of oocytes and their size as revealed using our statistical model might help to elucidate the mechanisms of dominant follicle selection and quality control of oocytes. Furthermore, for granulosa cells, we identified genes that were differentially expressed depending on their positions within the follicles, which suggests distinct epigenetic regulation and cell-cycle activities (Fig. 6). Thus, transcriptome analysis with DRaqL revealed a histology-associated, quantitative difference of transcriptomic profiles at the single-cell level.

In previous studies, direct application of the Smart-seq2 cDNA amplification, with non-denaturing cell lysis, has been employed for alcohol-fixed sections (Nichterwitz et al. 2016; Brasko et al. 2018; Deng et al. 2019; Lee et al. 2022). The lysis efficiency, which may depend on various parameters such as fixation conditions, has been controversial, and a purification-based approach has been proposed (Chen et al. 2017). In this study, our DRaqL-adapted methods achieved an improved cDNA amplification over the previous methods using non-denaturing cell lysis or RNA purification (Nichterwitz et al. 2016; Chen et al. 2017; Foley et al. 2019; Perez et al. 2021) (Fig. 4; Supplemental Figure S8).

The cDNA amplification methods used in this study were conducted in individual tubes, and thus their throughput would be comparable with those of previous studies using similar cDNA amplification approaches (e.g., the recent single-cell/low-input studies for primate gastrulae with Smart-seq2 (∼2,000 cells/samples) (Tyser et al. 2021; Bergmann et al. 2022)). In addition, the throughput could be further improved by using the multiplexed strategy employed in DRaqL-SMART-seq v4, DRaqL-Smart-seq2, and DRaqL-Protease-Smart-seq2.

On the other hand, the throughput might be limited by the one-by-one approaches for the LCM-based cell isolation, which are dependent on target tissues and the ease of identifying cells of interest. In this study, for the granulosa cells, the whole process of identifying target follicles and isolating cells of interest was manually conducted across many sections (typically, 1–2 target follicles per ovarian section met our investigation criteria). On the other hand, a high throughput, automated versatile LCM-based cell isolation method has been developed (Brasko et al. 2018) (>1,000 cells per day), which might improve the throughput of analysis.

In spite of the relatively low throughput of its cell isolation process, one of the advantages of LCM-based spatial transcriptomics over high-throughput methods for whole sections (see (Liao et al. 2021) for review) might be that it allows deeper transcriptomic analyses for any single cells of interest, including those of splice isoforms. A previous study aligned the cDNA sequences of ROIs to exons for investigation of the epi-transcriptome, but the quantitative performance for the expression levels of exon junctions remains elusive, with relatively low sensitivity of the method (a few thousand genes were detectable in slide-mounted culture cells in this previous report) (Lee et al. 2022). In this study, we performed quantitative exon–exon junction profiling of single cells with deep sequencing (Fig. 3) and detection of oocyte-specific splice isoforms (Supplemental Fig. S10), after 3’-end analyses with lower sequencing depths, thereby establishing a flexible single-cell experimental design for *in situ* transcriptomics. In addition, with respect to this particular application, our methods would have a significant advantage over a previous two-way LCM-combined method compatible with alcohol- and formalin-fixed sections (Foley et al. 2019), which relies on RNA hydrolysis and short-time PCR elongation, and thus restricts the analysis strictly to the 3’ ends of mRNA.

In conclusion, we have proposed an efficient, flexible analytical framework for single-cell transcriptomics from alcohol- and formalin-fixed tissue sections, which would serve as a complemental approach to the current high-throughput spatial and single-cell transcriptomics, as demonstrated with the discovery of histology-associated transcriptomic heterogeneity in the growing ovarian follicles.

## METHODS

The methods for library preparation, data processing, statistical modeling, animal sample preparation, cell culture, and the details of laser capture microdissection and cDNA amplification are described in the Supplemental Materials and Methods.

### Laser capture microdissection

Mouse ovaries and mESCs sunk in 10% polyvinyl alcohol (PVA) (Sigma Aldrich) were snap-frozen in liquid N_2_, and sectioned using a CM1860UV cryostat at a thickness of 15 μm, dried at room temperature, fixed and stained with 1% cresyl violet acetate, and washed with 100% isopropanol. Formalin fixation was performed with 10% formalin neutral buffer solution (Wako) at room temperature for 24 h. LCM was performed using the PALM MB4 laser microdissection system (Zeiss) with a ×20 or ×60 objective lens. Dissected cells were flicked into lysis buffer in caps of single, flat-top 200-μL PCR tubes (Greiner Bio-One). These procedures are summarized in Figure 1A.

### DRaqL-SC3-seq cDNA amplification for alcohol-fixed sections

Cells were isolated with LCM in 6.4 μL of cell lysis buffer (0.8 μL of GeneAmp 10xPCR Buffer II [Thermo Fisher], 0.48 μL of 25 mM MgCl_2_ [Thermo Fisher], 0.8 μL of 5% sodium deoxycholate [SDc] [Nacalai Tesque], 0.4 μL of 100 mM dithiothreitol [DTT] [Thermo Fisher], 0.64 μL of 40U/μL RNaseOUT [Thermo Fisher], 0.08 μL of 40U/μL porcine liver RNase inhibitor [Takara Bio], 0.16 μL of 2.5 mM dNTP [Takara Bio], 0.16 μL of 1:500,000 diluted ERCC RNA Spike-In Mix 1 [Thermo Fisher], 0.16 μL of 10 ng/μL V1[dT]_24_ primer [Hokkaido System Science], and 2.72 μL of deionized distilled water [DDW, Gibco]) in the flat caps of 0.2 mL PCR tubes (Greiner Bio-One), and spun down into the tubes by brief centrifugation. Cells were then lysed at 70°C for 6 min, followed by the addition of 2.8 μL quenching buffer (0.7 μL of Triton X-100 [Nacalai Tesque], 0.7 μL of 2% bovine serum albumin [BSA] [Takara Bio]) and incubation at 70°C for 90 sec to quench the denaturing effect of SDc. Subsequent cDNA synthesis and amplification were performed as described previously (Kurimoto et al. 2006). All oligonucleotides and thermal cycling programs used in this study are listed in Supplemental Table S1. DRaqL and adapted cDNA amplification are schematically represented in Figure 1A and 1B.

### DRaqL-SMART-Seq v4 cDNA amplification for alcohol-fixed sections

A Smart-seq v4 3’ DE Kit (Takara Bio) was adapted to DRaqL as follows. Cells were isolated with LCM in 6.4 μL of cell lysis buffer (0.25 μL of 40 U/μL Takara RNase Inhibitor, 0.4 μL of 100 mM DTT, 0.16 μL of 1:500,000 diluted ERCC RNA Spike-In Mix 1, 0.4 μL of 5% SDc, and 5.19 μL of nuclease-free water) in the caps of PCR tubes, and spun down into the tubes by brief centrifugation. Cells were then lysed at 72°C for 3 min, followed by the addition of 2.8 μL quenching buffer (0.7 μL of Triton X-100, 0.7 μL of 2% BSA, and 1.4 μL of 5× SuperScript II buffer [Thermo Fisher]) to reduce the denaturing effect of SDc. Then, 2.8 μL of RT oligo (1 μL of 12 μM Oligo dT In-line Primer and 1.8 uL of nuclease-free water: kit components) was added, followed by incubation at 72°C for 90 sec. Subsequent cDNA amplification was performed according to manufacturer’s instruction.

### DRaqL-Smart-seq2 cDNA amplification for alcohol-fixed sections

Cells were isolated with LCM in 6.4 μL of cell lysis buffer (0.6 μL of 5× SuperScript II buffer, 0.1 μL of 100 μM Oligo dT VN, 0.8 μL of dNTP mix [25 mM each], 0.25 μL of 40 U/μL recombinant RNase inhibitor [Takara Bio], 0.5 μL of 100 mM DTT, 0.06 μL of 1 M MgCl_2_ [Sigma-Aldrich], 2 μL of 5 M Betaine [Sigma-Aldrich], 0.16 μL of 1:500,000 diluted ERCC RNA Spike-In Mix 1, 0.4 μL of 5% SDc, and 1.53 μL of DDW) in the caps of PCR tubes. The isolated cells were lysed at 72°C for 6 min, followed by addition of 2.8 μL quenching buffer (0.7 μL of Triton X-100, 0.7 μL of 2% BSA, and 1.4 μL of 5× SuperScript II buffer [Thermo Fisher]). Then, 1.6 μL of a template-switching mixture (0.1 μL of 100 μM N-template-switching oligo, 0.25 μL of SuperScript II, 0.25 μL of SuperScript III, 0.2 μL of recombinant RNase inhibitor, and 0.8 μL of DDW) was added, followed by the cycling RT program for DRaqL-Smart-seq2. Then, cDNA amplification was performed by adding 15 μL of Seq amp PCR mixture (12.5 μL of 2x SeqAmp buffer (Takara Bio), 0.05 μL of 100 μM N-IS PCR primer, 0.5 μL of SeqAmp DNA polymerase (Takara Bio), and 1.95 μL of DDW) and applying the thermal cycling program for SeqAmp.

### DRaqL-Protease-Smart-seq2 cDNA amplification for formalin-fixed sections

Cells were isolated with LCM in 6.4 μL of cell lysis buffer (0.6 μL of 5× SuperScript II buffer, 0.1 μL of 100 μM Oligo dT VN, 0.8 μL of dNTP mix (25 mM each), 0.25 μL of 40 U/μL recombinant RNase inhibitor, 0.5 μL of 100 mM DTT, 0.06 μL of 1 M MgCl_2_, 2 μL of 5 M Betaine, 0.4 μL of 5% SDc, 0.32 μL of 900 mAU/mL Qiagen Protease, and 1.37 μL of DDW) in the caps of PCR tubes, and spun down into the tubes by brief centrifugation. The isolated cells were lysed by protease digestion at 50°C for 10 min followed by heat inactivation at 80°C for 15 min. The denaturing effect of SDc was quenched by addition of 0.8 μL of 1:2500,000 diluted ERCC RNA Spike-In Mix 1 and 2.8 μL of quenching buffer (0.7 μL of Triton X-100, 0.7 μL of 2% BSA, and 1.4 μL of 5× SuperScript II buffer [Thermo Fisher]), followed by incubation at 72°C for 90 sec. Then, 0.8 μL of a template-switching mixture (0.1 μL of 100 μM N-template-switching oligo, 0.25 μL of SuperScript II, 0.25 μL of SuperScript III, and 0.2 μL of recombinant RNase inhibitor) was added. The subsequent processes were performed as in DRaqL-Smart-seq2 cDNA amplification for alcohol-fixed sections described above.

## DATA ACCESS

All raw and processed sequencing data generated in this study have been submitted to the NCBI Gene Expression Omnibus (GEO; https://www.ncbi.nlm.nih.gov/geo/) under accession number GSE192551.

## ACKNOWLEDGEMENT

We thank all members of our laboratory for their discussion and advice regarding this study, and Junko Komeda for secretarial support. We also thank Kazusa Ohkita and the Single-CellGenome Information Analysis Core (SignAC) in ASHBi for the RNA sequence analysis. Finally, we thank Drs. Horvath and Migh for their kind advice about the experimental settings for LCM. This study was supported by KAKENHI grants (JP20H00471, J18K19295, and JP18H05553) to K.K; by funds from the Uehara Memorial Foundation; the Takeda Science Foundation; and the Daiichi Sankyo Foundation of Life Science to K.K; and by a KAKENHI grant (JP21K15107) to H.I.

## SUPPLEMENTAL DATA

The following are available in the Supplemental Data section:

**Supplemental Figure S1. Examination of DRaqL-SC3-seq cDNA amplification, related with Figure 1**

**Supplemental Figure S2. Evaluation of transcriptome data with DRaqL-SC3-seq, related with Figure 2**

**Supplemental Figure S3. Errors and biases of DRaqL-SC3-seq due to the use of sections, related with Figure 2**

**Supplemental Figure S4. Statistics of Y-adaptor sequencing, related with Figure 4**

**Supplemental Figure S5. DRaqL-SC3-seq of alcohol-fixed mouse ovarian sections, related with Figures 4 and 5**

**Supplemental Figure S6. DRaqL-SC3-seq of mouse oocytes from alcohol-fixed serial sections, related with Figure 5**

**Supplemental Figure S7. Bulk RNA-seq with DRaqL-SC3-seq of oocytes and granulosa cells in alcohol-fixed ovarian sections, related with Figure 4**

**Supplemental Figure S8. Adaptation of Smart-seq2-based cDNA amplification methods with DRaqL, and its application to formalin-fixed sections, related with Figure 4**

**Supplemental Figure S9. Morphological parameters of oocytes and their transcriptome, related with Figure 5**

**Supplemental Figure S10. Exon–exon junction analysis of mouse oocytes isolated from alcohol-fixed sections, related with Figure 6**

**Supplemental Figure S11. Previously published dataset of single-cell RNA-seq of oocytes and granulosa cells, related with Figure 6**.

**Supplemental Table S1. DNA primers and thermal cycling programs used in this study. Supplemental Table S2. Sequence statistics of mESCs**.

**Supplemental Table S3. Sequence statistics of oocytes and granulosa cells**.

**Supplemental Table S4. List of genes highly contributing to PC2**.

**Supplemental Table S5. List of genes highly correlated with oocyte diameters**.

**Supplemental Table S6. List of differentially expressed genes between granulosa cells neighboring and those not neighboring oocytes**.

**Supplemental Table S7. Histology and principal components of examined follicles**.

## REFFERENCES

Anderson E, Albertini DF. 1976. Gap junctions between the oocyte and companion follicle cells in the mammalian ovary. J Cell Biol 71: 680–686.

Bergmann S, Penfold CA, Slatery E, Siriwardena D, Drummer C, Clark S, Strawbridge SE, Kishimoto K, Vickers A, Tewary M et al. 2022. Spatial profiling of early primate gastrulation in utero. Nature 609: 136–143.

Brasko C, Smith K, Molnar C, Farago N, Hegedus L, Balind A, Balassa T, Szkalisity A, Sukosd F, Kocsis K et al. 2018. Intelligent image-based in situ single-cell isolation. Nat Commun 9: 226.

Byskov AG. 1974. Cell kinetic studies of follicular atresia in the mouse ovary. J Reprod Fertil 37: 277–285.

Chen J, Suo S, Tam PP, Han JJ, Peng G, Jing N. 2017. Spatial transcriptomic analysis of cryosectioned tissue samples with Geo-seq. Nat Protoc 12: 566–580.

Cheng R, Zheng X, Wang Y, Wang M, Zhou C, Liu J, Zhang Y, Quan F, Liu X. 2020. Genome-wide analysis of alternative splicing differences between oocyte and zygotedagger. Biol Reprod 102: 999–1010.

Deane HW. 1952. Histochemical observations on the ovary and oviduct of the albino rat during the estrous cycle. Am J Anat 91: 363–413.

Deng W, Xing C, David R, Mastroeni D, Ning M, Lo EH, Coleman PD. 2019. AmpliSeq Transcriptome of Laser Captured Neurons from Alzheimer Brain: Comparison of Single Cell Versus Neuron Pools. Aging Dis 10: 1146–1158.

Do DV, Strauss B, Cukuroglu E, Macaulay I, Wee KB, Hu TX, Igor RLM, Lee C, Harrison A, Butler R et al. 2018. SRSF3 maintains transcriptome integrity in oocytes by regulation of alternative splicing and transposable elements. Cell Discov 4: 33.

Eriksson T, Börjesson J, Tjerneld F. 2002. Mechanism of surfactant effect in enzymatic hydrolysis of lignocellulose. Enzyme and Microbial Technology 31: 353–364.

Espina V, Wulfkuhle JD, Calvert VS, VanMeter A, Zhou W, Coukos G, Geho DH, Petricoin EF, 3rd, Liotta LA. 2006. Laser-capture microdissection. Nat Protoc 1: 586–603.

Fan X, Moustakas I, Bialecka M, Del Valle JS, Overeem AW, Louwe LA, Pilgram GSK, van der Westerlaken LAJ, Mei H, Chuva de Sousa Lopes SM. 2021. Single-Cell Transcriptomics Analysis of Human Small Antral Follicles. Int J Mol Sci 22.

Foley JW, Zhu C, Jolivet P, Zhu SX, Lu P, Meaney MJ, West RB. 2019. Gene expression profiling of single cells from archival tissue with laser-capture microdissection and Smart-3SEQ. Genome Res 29: 1816–1825.

Ghimire S, Stewart CG, Thurman AL, Pezzulo AA. 2021. Performance of a scalable RNA extraction-free transcriptome profiling method for adherent cultured human cells. Sci Rep 11: 19438.

Goldenberger D, Perschil I, Ritzler M, Altwegg M. 1995. A simple “universal” DNA extraction procedure using SDS and proteinase K is compatible with direct PCR amplification. PCR Methods Appl 4: 368–370.

Gu C, Liu S, Wu Q, Zhang L, Guo F. 2019. Integrative single-cell analysis of transcriptome, DNA methylome and chromatin accessibility in mouse oocytes. Cell Res 29: 110–123.

Hamazaki N, Kyogoku H, Araki H, Miura F, Horikawa C, Hamada N, Shimamoto S, Hikabe O, Nakashima K, Kitajima TS et al. 2021. Reconstitution of the oocyte transcriptional network with transcription factors. Nature 589: 264–269.

Jonstromer M, Strey R. 1992. Nonionic bilayers in dilute-solutions—effect of additives. J Phys Chem 96: 5993–6000.

Kasowitz SD, Ma J, Anderson SJ, Leu NA, Xu Y, Gregory BD, Schultz RM, Wang PJ. 2018. Nuclear m6A reader YTHDC1 regulates alternative polyadenylation and splicing during mouse oocyte development. PLoS Genet 14: e1007412.

Kurimoto K, Yabuta Y, Ohinata Y, Ono Y, Uno KD, Yamada RG, Ueda HR, Saitou M. 2006. An improved single-cell cDNA amplification method for efficient high-density oligonucleotide microarray analysis. Nucleic Acids Res 34: e42.

Kurimoto K, Yabuta Y, Ohinata Y, Shigeta M, Yamanaka K, Saitou M. 2008. Complex genome-wide transcription dynamics orchestrated by Blimp1 for the specification of the germ cell lineage in mice. Genes Dev 22: 1617–1635.

Le AV, Huang D, Blick T, Thompson EW, Dobrovic A. 2015. An optimised direct lysis method for gene expression studies on low cell numbers. Sci Rep 5: 12859.

Lee AC, Lee Y, Choi A, Lee HB, Shin K, Lee H, Kim JY, Ryu HS, Kim HS, Ryu SY et al. 2022. Spatial epitranscriptomics reveals A-to-I editome specific to cancer stem cell microniches. Nat Commun 13: 2540.

Li J, Lu M, Zhang P, Hou E, Li T, Liu X, Xu X, Wang Z, Fan Y, Zhen X et al. 2020. Aberrant spliceosome expression and altered alternative splicing events correlate with maturation deficiency in human oocytes. Cell Cycle 19: 2182–2194.

Li R, Albertini DF. 2013. The road to maturation: somatic cell interaction and self-organization of the mammalian oocyte. Nat Rev Mol Cell Biol 14: 141–152.

Li S, Chen LN, Zhu HJ, Feng X, Xie FY, Luo SM, Ou XH, Ma JY. 2021. Single-cell RNA sequencing analysis of mouse follicular somatic cells. Biol Reprod doi:10.1093/biolre/ioab163.

Liao J, Lu X, Shao X, Zhu L, Fan X. 2021. Uncovering an Organ’s Molecular Architecture at Single-Cell Resolution by Spatially Resolved Transcriptomics. Trends Biotechnol 39: 43–58.

Lieberman-Aiden E, van Berkum NL, Williams L, Imakaev M, Ragoczy T, Telling A, Amit I, Lajoie BR, Sabo PJ, Dorschner MO et al. 2009. Comprehensive mapping of long-range interactions reveals folding principles of the human genome. Science 326: 289–293.

Marks H, Kalkan T, Menafra R, Denissov S, Jones K, Hofemeister H, Nichols J, Kranz A, Stewart AF, Smith A et al. 2012. The transcriptional and epigenomic foundations of ground state pluripotency. Cell 149: 590–604.

Matzuk MM, Burns KH, Viveiros MM, Eppig JJ. 2002. Intercellular communication in the mammalian ovary: oocytes carry the conversation. Science 296: 2178–2180.

Nakamura T, Okamoto I, Sasaki K, Yabuta Y, Iwatani C, Tsuchiya H, Seita Y, Nakamura S, Yamamoto T, Saitou M. 2016. A developmental coordinate of pluripotency among mice, monkeys and humans. Nature 537: 57–62.

Nakamura T, Yabuta Y, Okamoto I, Aramaki S, Yokobayashi S, Kurimoto K, Sekiguchi K, Nakagawa M, Yamamoto T, Saitou M. 2015. SC3-seq: a method for highly parallel and quantitative measurement of single-cell gene expression. Nucleic Acids Res 43: e60.

Nichterwitz S, Chen G, Aguila Benitez J, Yilmaz M, Storvall H, Cao M, Sandberg R, Deng Q, Hedlund E. 2016. Laser capture microscopy coupled with Smart-seq2 for precise spatial transcriptomic profiling. Nat Commun 7: 12139.

Paavilainen L, Edvinsson A, Asplund A, Hober S, Kampf C, Ponten F, Wester K. 2010. The impact of tissue fixatives on morphology and antibody-based protein profiling in tissues and cells. J Histochem Cytochem 58: 237–246.

Pan H, O’Brien M J, Wigglesworth K, Eppig JJ, Schultz RM. 2005. Transcript profiling during mouse oocyte development and the effect of gonadotropin priming and development in vitro. Dev Biol 286: 493–506.

Perez JD, Dieck ST, Alvarez-Castelao B, Tushev G, Chan IC, Schuman EM. 2021. Subcellular sequencing of single neurons reveals the dendritic transcriptome of GABAergic interneurons. Elife 10.

Picelli S, Bjorklund AK, Faridani OR, Sagasser S, Winberg G, Sandberg R. 2013. Smart-seq2 for sensitive full-length transcriptome profiling in single cells. Nat Methods 10: 1096–1098.

Simon AM, Goodenough DA, Li E, Paul DL. 1997. Female infertility in mice lacking connexin 37. Nature 385: 525–529.

Sivars U, Bergfeldt K, Piculell L, Tjerneld F. 1994. Effect of ionic surfactants on nonionic bilayers—bending elasticity of weakly charged membrane. J Phys Chem 98: 3908–3912.

Svensson V, Vento-Tormo R, Teichmann SA. 2018. Exponential scaling of single-cell RNA-seq in the past decade. Nat Protoc 13: 599–604.

Tang F, Barbacioru C, Wang Y, Nordman E, Lee C, Xu N, Wang X, Bodeau J, Tuch BB, Siddiqui A et al. 2009. mRNA-Seq whole-transcriptome analysis of a single cell. Nat Methods 6: 377–382.

Titford ME, Horenstein MG. 2005. Histomorphologic assessment of formalin substitute fixatives for diagnostic surgical pathology. Arch Pathol Lab Med 129: 502–506.

Tyser RCV, Mahammadov E, Nakanoh S, Vallier L, Scialdone A, Srinivas S. 2021. Single-cell transcriptomic characterization of a gastrulating human embryo. Nature doi:10.1038/s41586-021-04158-y.

Waylen LN, Nim HT, Martelotto LG, Ramialison M. 2020. From whole-mount to single-cell spatial assessment of gene expression in 3D. Commun Biol 3: 602.

Williams CJ, Erickson GF. 2000. Morphology and Physiology of the Ovary. In Endotext, (ed. KR Feingold, et al.), South Dartmouth (MA).

Ying QL, Wray J, Nichols J, Batlle-Morera L, Doble B, Woodgett J, Cohen P, Smith A. 2008. The ground state of embryonic stem cell self-renewal. Nature 453: 519–523.

Yu L, Zhang H, Guan X, Qin D, Zhou J, Wu X. 2021. Loss of ESRP1 blocks mouse oocyte development and leads to female infertility. Development 148.

Zhang Y, Yan Z, Qin Q, Nisenblat V, Chang HM, Yu Y, Wang T, Lu C, Yang M, Yang S et al. 2018. Transcriptome Landscape of Human Folliculogenesis Reveals Oocyte and Granulosa Cell Interactions. Mol Cell doi:10.1016/j.molcel.2018.10.029.

